# NEUTROPHILS ARE INDISPENSABLE FOR ADVERSE CARDIAC REMODELING IN HEART FAILURE

**DOI:** 10.1101/2023.10.31.565035

**Authors:** Sergey Antipenko, Nicolas Mayfield, Miki Jinno, Matthias Gunzer, Mohamed Ameen Ismahil, Tariq Hamid, Sumanth D. Prabhu, Gregg Rokosh

## Abstract

Persistent immune activation contributes significantly to left ventricular (LV) dysfunction and adverse remodeling in heart failure (HF). In contrast to their well-known essential role in acute myocardial infarction (MI) as first responders that clear dead cells and facilitate subsequent reparative macrophage polarization, the role of neutrophils in the pathobiology of chronic ischemic HF is poorly defined. To determine the importance of neutrophils in the progression of ischemic cardiomyopathy, we measured their production, levels, and activation in a mouse model chronic HF 8 weeks after permanent coronary artery ligation and large MI. In HF mice, neutrophils were expanded both locally in failing myocardium (more in the border zone) and systemically in the blood, spleen and bone marrow, together with increased BM granulopoiesis. There were heightened stimuli for neutrophil recruitment and trafficking in HF, with increased myocardial expression of the neutrophil chemoattract chemokines CXCL1 and CXCL5, and increased neutrophil chemotactic factors in the circulation. HF neutrophil NETotic activity was increased in vitro with coordinate increases in circulating neutrophil extracellular traps (NETs) in vivo. Neutrophil depletion with either antibody-based or genetic approaches abrogated the progression of LV remodeling and fibrosis at both intermediate and late stages of HF. Moreover, analogous to murine HF, the plasma milieu in human acute decompensated HF strongly promoted neutrophil trafficking. Collectively, these results support a key tissue-injurious role for neutrophils and their associated cytotoxic products in ischemic cardiomyopathy and suggest that neutrophils are potential targets for therapeutic immunomodulation in this disease.

## INTRODUCTION

Chronic ischemic heart failure (HF) is characterized by local and systemic expansion of innate and adaptive immune cells that promote adverse left ventricular (LV) remodeling.^1–4^ Globally, augmented myelopoiesis driven by factors such as sympathetic activation, circulating cytokines, and growth factors^3, 5^ generates a pathological milieu for myeloid cell activation. Indeed, several studies have indicated critical roles for pro-inflammatory monocytes and tissue macrophages in the progression of LV remodeling after myocardial infarction (MI).^6–9^ In contrast, surprisingly little is known about the role of neutrophils in chronic ischemic cardiomyopathy. Neutrophils are the first leukocytes to infiltrate the heart in response to acute MI and are essential for proper wound healing, providing phagocytic clearance of dead cells and facilitating reparative macrophage polarization and subsequent resolution of inflammation.^10, 11^ Sex-, age-, and time-of-day-related differences in neutrophil phenotype may underlie differences in outcomes after acute MI.^12, 13^ Beyond *acute* post-MI tissue repair and wound healing, however, there is little understanding as to the importance of neutrophils in the pathogenesis of *chronic* LV remodeling.

While neutrophils are essential for post-MI wound healing, prolonged neutrophil activity and persistence can exacerbate myocardial injury due to their cytotoxic effector properties.^14^ These include phagocytosis and release of granule components (e.g., proteases, oxidants, antimicrobial peptides) into the phagosome or extracellular space, free radical production, and the formation of neutrophil extracellular traps (NETs). NETs are large 3-D structures composed of thin chromatin DNA associated with histones, granular enzymes, and peptides expressed during NETosis, a form of regulated neutrophil cell death.^15, 16^ The release of granular contents and NETs, and apoptotic release of lysophosphatidylcholine, increases monocyte recruitment and activation.^17, 18^ In a transverse-aortic constriction (TAC) mouse model, cardiac neutrophil expansion occurred early (3–7 d) after pressure-overload, prior to the development of HF, and did not persist at later stages.^19^ Neutrophil depletion prior to TAC and myeloid loss of Wnt5a ameliorated pressure-overload cardiac remodeling. Similarly, cardiac neutrophil expansion was observed early after angiotensin II infusion in mice, again without persistence at later stages and prior to the development of LV remodeling, with an integral role for myeloid KLF2 in remodeling progression.^20^ Notably, the immune cell activation profile after MI, and during the progression of post-MI LV remodeling, differs considerably from non-ischemic HF.^1, 11^ Accordingly, here we sought to define the neutrophil profile and the function of neutrophil effectors in chronic ischemic cardiomyopathy, and determine their pathophysiological role in adverse LV remodeling.

## METHODS

### Mouse models

All animal studies were approved by the Institutional Animal Care and Use Committee of the University of Alabama at Birmingham or Washington University in St. Louis and were compliant with the National Institutes of Health Guide for the Care and Use of Laboratory Animals (DHHS publication No. 85-23, revised 1996). A total of 296 mice were used. C57BL/6J wild-type (WT) and ROSA26iDTR (iDTR) mice were obtained from Jackson Laboratories (stock #000664 and #007900, respectively), while Ly6G-Cre (Catchup) mice were developed by Dr. Mathias Gunzer (University Duisburg–Essen) as previously described.^21^ Ly6G-DTR mice were generated by crossing Catchup and iDTR mice.

### *In vivo* experimental protocols

Under 1-2% inhaled isoflurane anesthesia (in 100% O2) and mechanical ventilation, 10-12 week old male mice underwent permanent left coronary artery ligation surgery to induce MI and ischemic HF or sham operation as previously described.^2, 4, 22, 23^ For different treatments, HF mice with comparable degrees of adverse LV remodeling by echocardiography at 4- or 8-weeks post-MI were randomized to different treatment groups. Interventions were rigorously performed at prespecified times each day given the known circadian rhythmicity of circulating neutrophil levels.^13^ For antibody-mediated neutrophil ablation, WT mice received anti-Ly6G (clone 1A8) or IgG isotype control (BioXcell BE0075-1 and BE0089, respectively) at a dose of 250 μg/mouse i.p. every 4 days at zeitgeber time 5 (ZT5) beginning at either 28 and 56 d post-surgery, for a total of 28 d (8 total doses). Neutrophil ablation in Ly6G-DTR mice was induced 56 d post-MI via i.p. diphtheria toxin (DT; 10 μg/kg/d; Sigma D0564) administered twice daily at ZT0 and ZT12 for 2 w, with PBS used as a vehicle control.

### Echocardiography

Two-dimensional echocardiography was performed on anesthetized mice (1-2% inhaled isoflurane) and continuous ECG monitoring using a VisualSonics Vevo770 or Vevo3100 High-Resolution System and 30 MHz RMV707B or MX400 transducer with a heated, bench-mounted adjustable rail system.^2, 4, 22, 24^ Body temperature was maintained at 37.0 ± 0.5°C and heart rate at 500 ± 50 bpm. Images were taken in the parasternal long-axis view and LV chamber volumes in end-diastole (EDV) and end-systole (ESV) were determined using the area-length method from images captured in ECG-gated Kilohertz Visualization mode. LV ejection fraction (EF) was calculated as [(EDV−ESV)/EDV] *100.

### Immune cell isolation and flow cytometry

All cell and tissue collections were performed between ZT3 and ZT7. Live mononuclear cells were isolated from the blood, heart, spleen, and bone marrow and processed for flow cytometry as previously described.^2, 4, 22, 24, 25^ Peripheral blood (∼100 μL) was collected via cheek vein and placed in EDTA coated tubes (BD Biosciences 365974). Erythrocytes were lysed using RBC Lysis Buffer (eBioscience, ThermoFisher Scientific 00-4333-57) before washing with PBS. Whole hearts were flushed with PBS, minced into small pieces, suspended in RPMI 1640 medium (Gibco, ThermoFisher Scientific 11875119) and spun at low speed (50*g*) for 5 min at 4°C to remove circulating cells and then digested in RPMI 1640 with 1 mg/mL Type 2 collagenase (Worthington Biochemical LS004177) for 45 min at 37°C, agitating the tissue mixture every 5 min. Cell suspensions were filtered through a 40-μm cell strainer into RPMI with 10% FBS (GE Healthcare Life Sciences SH30071.03) and incubated for 5 min at 37°C. Isolates were centrifuged (500*g* for 5 min) before collecting cells and washing with PBS. A portion of spleen was taken and manually homogenized to collect splenocytes. To collect bone marrow cells, tibias were clipped at the ends of the bones and flushed with autoMACS Running Buffer (Miltenyi Biotec 130-091-221) using a 22G syringe.

Once processed, all cells were fixed with 1% paraformaldehyde on ice for 10 minutes for later flow cytometric analysis, followed by washing with PBS and resuspending in autoMACS Running Buffer. As appropriate for specific protocols, cells were stained for membrane markers by incubating with antibodies against CD45-BV605 (ThermoFisher Scientific 83-0451-42), Ly6G-PE (ThermoFisher Scientific 12-5931-82), CD11b-AF700 (ThermoFisher Scientific 56-0112-82), Lineage cocktail-Pacific Blue (Biolegend 133310), Sca1-PerCP (Biolegend 108122), cKit-PECy7 (Biolegend 135112), CD127-BV605 (Biolegend 135041), CD34-FITC (ThermoFisher Scientific 11-0341-81), CD16/32-PE (Biolegend 133310) for 1 h. All staining was performed on ice to maintain cell morphology. Flow cytometric data were acquired using either a BD Biosciences LSR II or an Invitrogen Attune NXT flow cytometer and analyzed using FlowJo software v10.6.1.

### Assessment of NETs and NETosis

Peripheral blood was isolated, and erythrocytes were lysed as described for flow cytometry. Neutrophils were isolated by negative selection using the Mojosort Mouse Neutrophil Isolation Kit (Biolegend 480058). Wells were pre-coated with 100 μL per well of capture anti-histone antibody contained in the Cell Death Detection ELISA kit (1:40 in 1x coating buffer; Roche 11544675001) or anti-myeloperoxidase (MPO, Upstate 07-496) (1:10) by incubating at room temperature (RT) for 1 h followed by 200 μL incubation buffer from the Cell Death Detection ELISA kit for 30 min at RT. Incubation buffer was removed and plates were washed three times with 250 μL of Wash Solution from the Cell Death Detection ELISA kit before seeding 50,000 neutrophils per well in 100 μL RPMI media containing 1% FBS (or 20 μL plasma with 80 μL with incubation buffer per well) incubating at 37°C for 1 h. In each well, 40 μL of DNase I (Roche 04716728001, 1:1000 in DNase I dilution buffer) was added and incubated at RT for 15 minutes while shaking at 250 rpm. This reaction was neutralized with 10 μL of 25 mM EGTA (2.5 mM final concentration) at 4°C overnight. Plates were washed three times with 250 μL ELISA wash solution before adding 100 μL ELISA anti-DNA-HRP and incubating at RT for 90 min. Finally, plates were washed three times with 250 μL Wash Solution and incubated at RT for 15 min while shaking at 250 rpm in 100 μL of ELISA substrate solution. This colorimetric assay for NETs was measured at an absorbance of 405 nm using a Biotek plate reader.

### Neutrophil chemotaxis and transmigration

Peripheral blood was isolated from naive mice, erythrocytes were lysed as described above. Neutrophils were then sorted by negative selection using Mojosort Mouse Neutrophil Isolation Kit (Biolegend 480058) and stained with 100 nM dichlorofluorescein diacetate (DCFDA, ThermoFisher Scientific D399) for visualization. Neutrophils were seeded on a fibronectin coated 8-chamber slide at a concentration of 50,000 cells/100 μL RPMI with 1% FBS. A chemotactic gradient was created by loading 10 μL of plasma into the tip of a patch clamp micropipette and placing it along the edge of a chamber. The chemotactic potential of the plasma was determined by measuring neutrophil migration properties (line speed, line length, and straightness) via time-lapse confocal microscopy (Nikon C2+ Confocal Microscope) over a 3-minute period using Nikon Elements AR software v4.30.01.

Transmigration towards chemoattractants CXCL1 (100 ng/mL), CXCL5 (200 ng/mL), granulocyte colony stimulating factor (G-CSF, 100 ng/mL), mouse formylated peptide from NADH:ubiquinone oxidoreductase chain 6 f-MNNYIF (f-MNN, 100nM), mouse mitochondrial non-formylated peptide control (MNN, 100nM), CXCL8 (100 ng/mL), CCL3 (100 ng/mL), human formylated peptide from NADH-ubiquinone oxidoreductase chain 6 f-MMYALF (f-MMY, 100 nM), human non-formylated peptide control (MMY, 100 nM) was determined by measuring transmigration of isolated neutrophils (5 x 10^4^ cells/well) through a 3 μm Transwell membrane upon 2 h of incubation with the chemoattractant of interest at 37°C. Transmigration reactions were stopped with addition of 0.5 mM EDTA and migrated cells were counted using a hemacytometer and expressed as a percent of total cells.

### Human studies

Peripheral blood was obtained from human subjects hospitalized with acute decompensated HF (ADHF) (UAB IRB 300000114) and ambulatory healthy controls. Blood was collected in EDTA-treated tubes and plasma was obtained upon centrifugation for 15 min at 2,000*g* and then stored at -80°C for future use. Human blood neutrophils from healthy controls were isolated using negative sorting with the EasySep Direct Human Neutrophil Isolation Kit (STEMCELL Technologies 19666), loaded with DCFDA, and used within 4 hours for studies of chemotaxis (using time-lapse confocal microscopy) and transmigration (using Transwell chambers), as described above.

### Histological analysis

Fibrosis was measured using Masson’s Trichrome staining, as previously described.^2, 4, 23, 26–29^ Fibrosis was quantified in the LV border and remote zones using Metamorph software v6.3r5 (Molecular Devices). Consistent collagen (blue) and cardiomyocyte (red) staining thresholds were maintained while all fibrotic areas were captured. Tissue fibrosis was determined from 4-6 high-power fields per section and expressed as a percent of total cross-sectional area. For cardiac neutrophil quantitation, LV sections were paraffin embedded, rehydrated, and stained with rat anti-mouse Ly6G (1:50, Biolegend 127602) with secondary goat anti-rat Alexa Fluor 488 (1:100, Invitrogen A-11008). Nuclei were stained with DAPI. Ly6G^+^ cells were counted in five high-power fields (60x) per LV region (remote and border zone). Images were acquired using a Nikon A1 confocal microscope and Nikon Elements AR software v4.30.01.

### RT-qPCR

RNA extraction from LV tissue (encompassing remote and border zone myocardium while excluding infarct tissue), cDNA synthesis, and quantitative real-time PCR were performed as previously described.^2, 4, 23, 26–29^ Gene expression was determined for CXCL1, CXCL5, CCL17, G-CSF, granulocyte-macrophage stimulating factor (GM-CSF), and macrophage stimulating factor (M-CSF). Forward and reverse primer pairs used to determine gene transcript levels are provided in *Table 1*. Gene expression was normalized to β-actin for LV tissue using the ΔΔCT comparative method and expressed as fold-change.

**Table 1.**
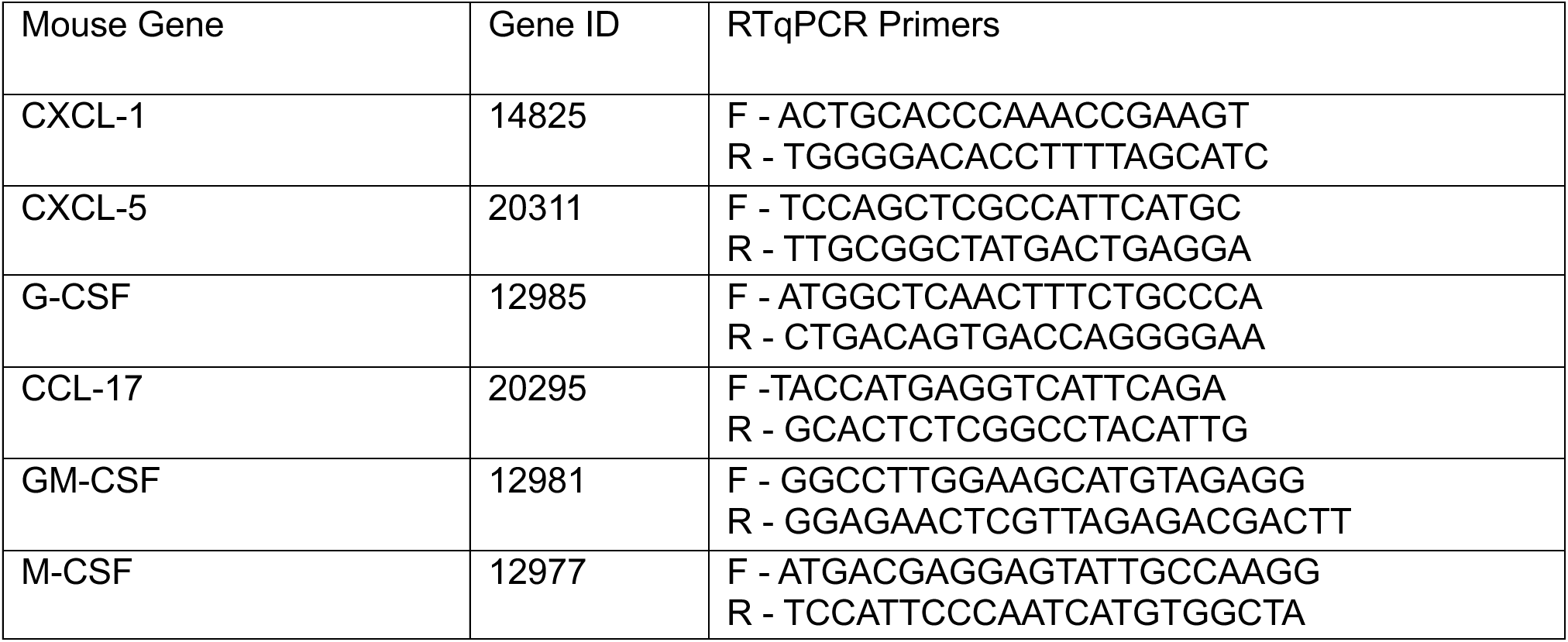
RTqPCR primers for gene expression in failing hearts.

### Statistical Analysis

All summative data are shown as box plots with representations of 25^th^ percentile, median and 75^th^ percentile. Group variances were compared using the Brown−Forsythe test, while normality was assessed using the D’Agostino–Pearson test. Statistical comparison between two groups was performed using two-tailed paired or unpaired t-tests with equal or unequal variance, as appropriate. Non-normal distributions were assessed using the Mann-Whitney test for non-parametric analyses. Comparisons of more than two groups used a one-or two-way ANOVA with Dunnett’s post-test (for normal distribution) or Kruskal-Wallis test with Dunn’s post-test (for non-normal distribution). The specific approaches applied are stated in the figure legends. All analyses were conducted using GraphPad Prism version 8.4.1. A p < 0.05 was considered significant.

## RESULTS

### Neutrophils expand locally and systemically in chronic ischemic HF

Adult male C57Bl/6 mice underwent permanent coronary ligation or sham operation. Large infarctions were confirmed by echocardiography 2 d after MI, and mice were assessed at 4 and 8 w post-MI (*Figure 1A)*. At 8 w post-MI, as compared to sham-operated mice, ligated mice exhibited hallmarks of ischemic cardiomyopathy and HF, with significantly increased LV end-diastolic volume (EDV) and end-systolic volume (ESV), reduced LV ejection fraction (EF), increased heart, lung, and spleen weight, and border and remote zone fibrosis (*<u>Figure S1</u>*). CD45^+^ leukocytes were isolated from the heart, peripheral blood, spleen, and bone marrow (BM), and CD11b^+^Ly6G^+^ neutrophils were identified using gating strategies depicted in *Figure 1B* and *<u>Figure S2</u>*. Cardiac neutrophils were expanded nearly 3-fold in the failing heart (*Figure 1B*), accompanied by significantly increased neutrophils in the circulation, spleen, and BM (*Figure 1C*). Cardiac immunostaining (*Figure 1D*) indicated that neutrophils were localized in the interstitium of both the MI border zone and zones remote from the infarct, with particularly striking spatial expansion (∼25-fold) in the border zone.

**Figure 1.**
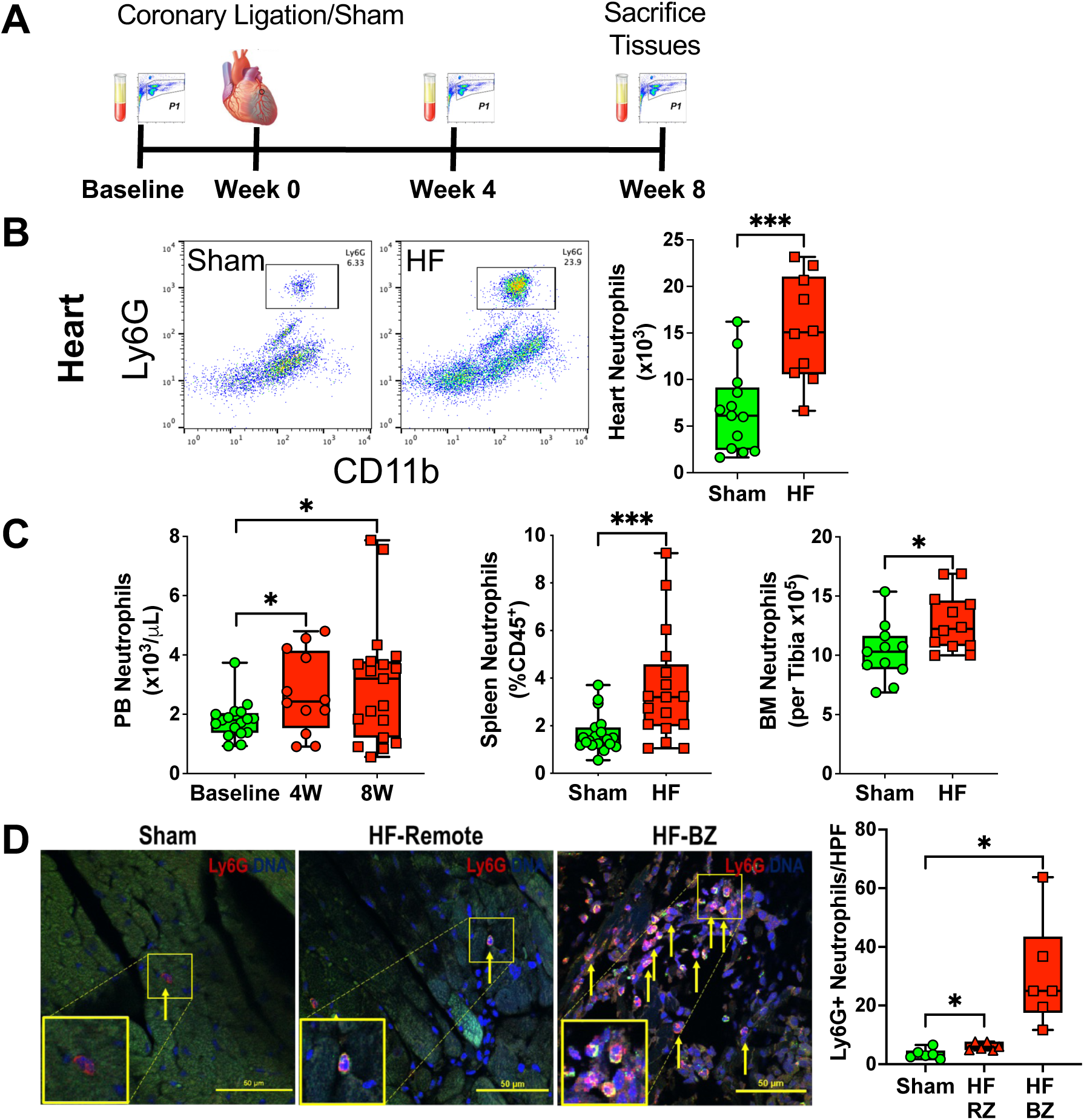
Neutrophil expansion in HF mice. ***A***, Neutrophil levels were quantified by flow cytometry (CD45^+^CD11b^+^Ly6G^+^ cells) in heart, bone marrow, peripheral blood (PB), and spleen from HF and sham mice at baseline (for PB) and 4 and/or 8 w post-MI or sham surgery. ***B***, Representative flow plots and corresponding quantitation of neutrophils in sham-operated (n=13) and failing mouse hearts (n=10) 8 w post-MI or sham surgery. ***C***, PB neutrophils at baseline, 4 w and 8 w post-MI in HF mice, and splenic and bone marrow neutrophils in sham and HF mice 8 w post-MI or sham surgery (n as indicated in panels). ***D***, Immunofluorescent staining for Ly6G^+^ neutrophils in heart sections from sham and HF mice (8 w post-MI or sham surgery) and corresponding quantitation; RZ, remote zone, BZ, MI border zone. DAPI (DNA) Blue, Ly6G Red; scale bar = 50 µm. The average number of neutrophils per high-powered field (HPF) were determined from 5 areas per section. Sham n=6, HF RZ n=6, HF BZ n=6. *p≤0.05, *** p≤0.001 versus sham by two-tailed unpaired t-test (***B*** and ***C***) or one-way ANOVA with Dunnett’s post-test (***D***).

As neutrophils are short-lived cells with a lifespan of 6-48 h,^30^ they are constantly produced by granulopoiesis in the BM (up to 10^11^ cells/day). Hence, augmented granulopoiesis would be obligatory for neutrophil expansion to ensure population replenishment. Accordingly, we examined hematopoietic stem and granulocyte progenitor cell populations in the BM of sham and HF mice (8 w post-MI) by flow cytometry (*<u>Figure S2</u>*). As illustrated in *Figure 2*, multipotent myeloid progenitors (Lineage^−^Sca1^−^cKit^+^) progenitors and CD127^−^CD34^+^CD16/32^+^ granulocyte-monocyte progenitors (GMPs) within this population^31, 32^ were significantly increased in HF mice. Lineage^−^ Sca1^−^cKit^+^CD127^−^CD34^+^CD16/32^−^ common myeloid progenitors (CMPs) and Lin^−^Sca1^+^cKit^+^ (LSK) hematopoietic stem cells tended to increase in HF mice (p<0.07 and 0.09 versus sham, respectively). Hence, HF mice exhibited local and systemic neutrophilia in concert with increased BM granulopoiesis.

**Figure 2.**
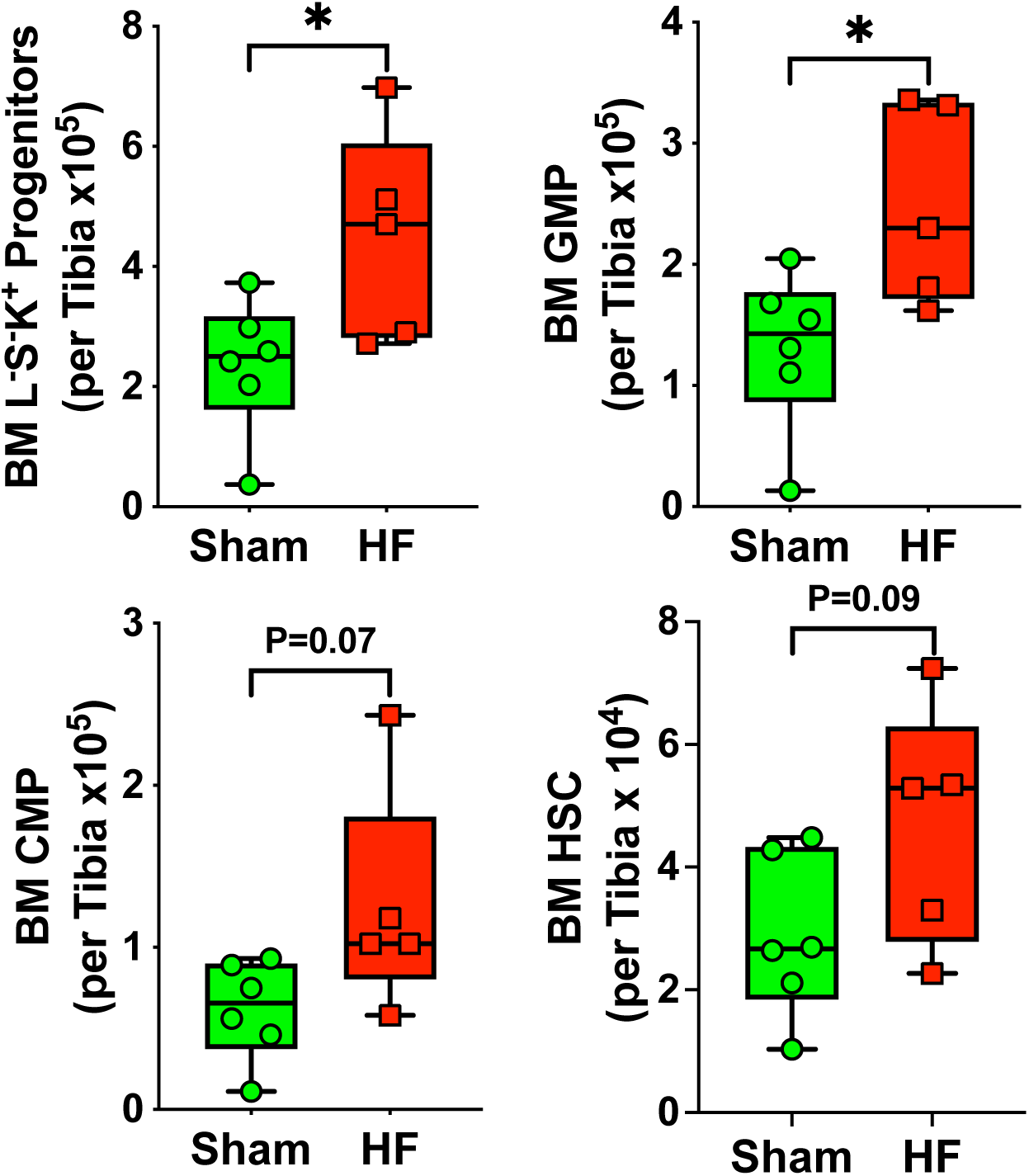
Increased granulopoiesis in HF. Granulopoiesis in sham-operated and heart failure (HF) mice was evaluated by determining bone marrow progenitor and stem cell numbers by flow cytometry at 8 w post-MI or sham surgery. L^-^S^-^K^+^, Lineage^-^Sca1^-^Kit^+^ multipotent myeloid progenitors; GMP, granulocyte monocyte progenitors (Lineage^-^Sca1^-^cKit^+^CD127^-^ CD34^+^CD16/32^+^); CMP, common myeloid progenitors (Lineage^-^Sca1^-^cKit^+^CD127^-^ CD34^+^CD16/32^-^); HSC, LSK hematopoietic stem cells (Lineage^-^Sca1^+^cKit^+^). Sham n=6, HF n=5. *p≤0.05 versus sham by two-tailed unpaired t-test.

### Augmented stimuli for neutrophil recruitment in murine HF

Neutrophil expansion in HF suggests a microenvironment supporting neutrophil recruitment. Neutrophils are attracted to injured tissue by a broad range of chemoattractants, including damage associated molecular patterns (DAMPs) and CXC chemokines containing the glutamic acid-leucine-arginine (ELR) motif.^11, 33^ As shown in *Figure 3*, real time RT-PCR revealed significantly increased gene expression of the neutrophil chemoattractants CXCL1 and CXCL5 in failing hearts versus sham hearts. Moreover, expression of CCL17, a T-cell chemoattractant that can be secreted by neutrophils,^34^ was also increased in failing hearts. In contrast, there was no change in cardiac expression of myeloid differentiation factors (G-CSF, GM-CSF, and M-CSF) in HF. Increased CXCL1 and CXCL5 expression provides a potential stimulus to recruit neutrophils to the heart via activation of their cognate receptor CXCR2. To determine whether circulating chemotactic factors are increased in HF, plasma was collected from naïve and HF mice and used to stimulate chemotaxis of labeled naïve neutrophils *in vitro*, as quantitated by time-lapse confocal imaging and vector analysis. As compared with plasma from naïve mice (or no plasma control), HF plasma induced significantly greater neutrophil migration, evidenced by with increased cell distance traveled (line length), rate of migration toward the plasma source (line speed), and increased directedness toward the plasma source (*Figures 4A-C*). Moreover, in Transwell chemotaxis assays, CXCL1 and CXCL5 significantly increased naïve neutrophil chemotaxis (*Figure 4D*). Neutrophils similarly responded to mitochondrial formyl methionine-asparagine-asparagine (fMNN), an important DAMP released upon necrotic cell death (but not to MNN alone).^35^

**Figure 3.**
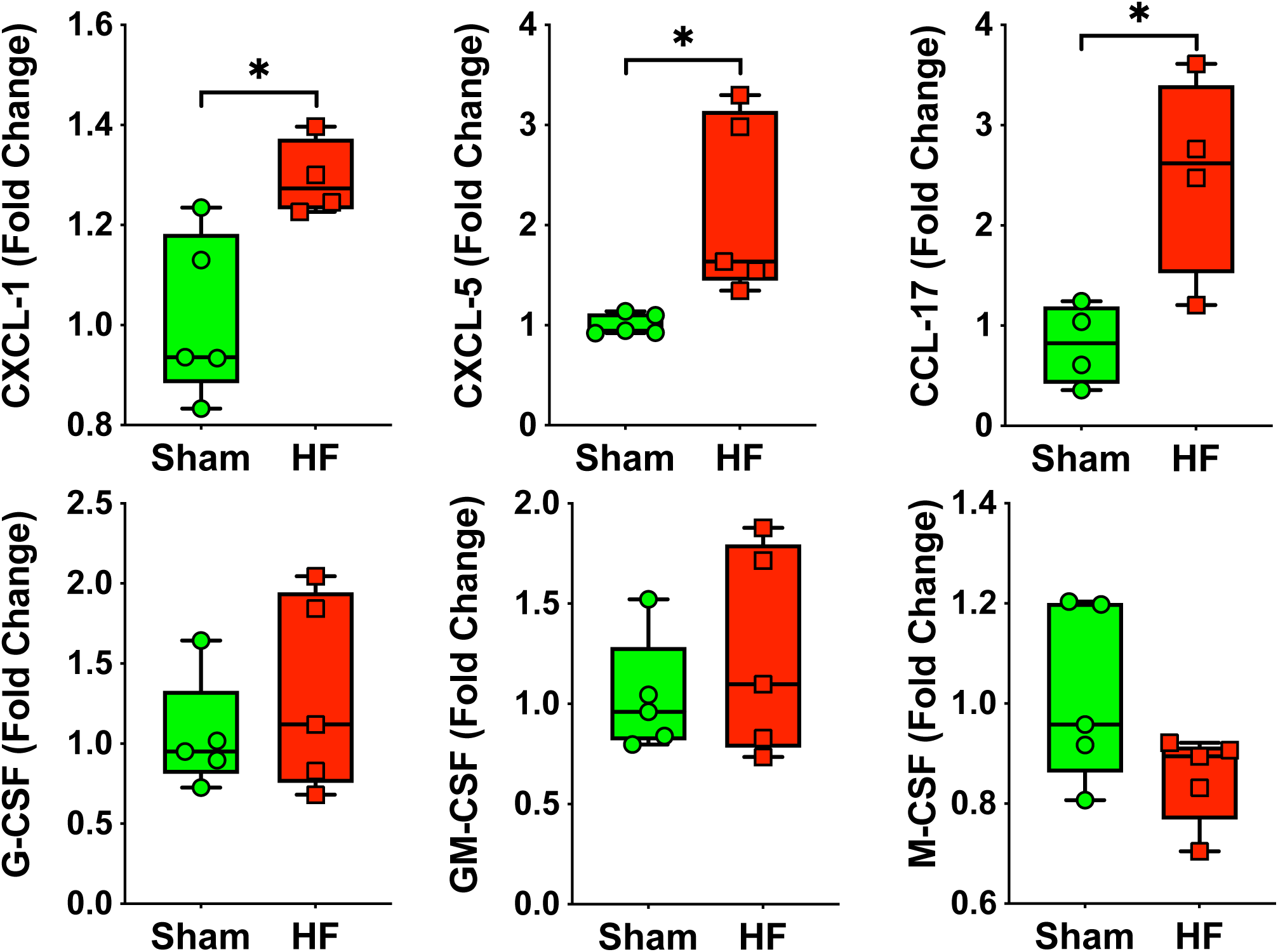
Chemokine and myeloid differentiation factor expression in the failing heart. Gene expression by real time RT-PCR of CXCL1 and CXCL5 (neutrophil chemoattractants), CCL17, and myeloid differentiation factors G-CSF, GM-CSF, and M-CSF in sham-operated and heart failure (HF) mice 8 w after surgery. N=4-5 per group. *p≤0.05 versus sham by two-tailed unpaired t-test.

**Figure 4.**
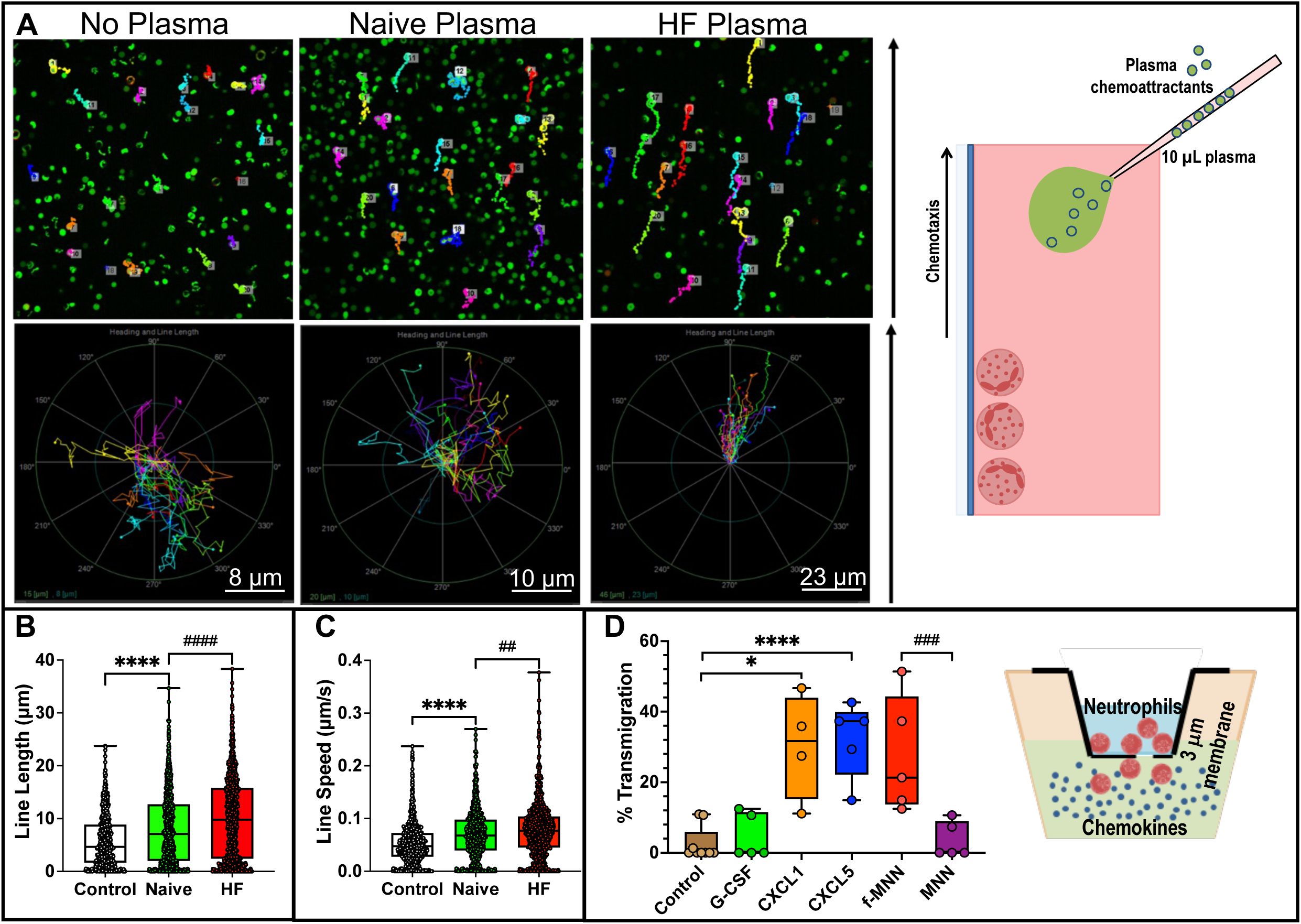
Increased stimuli for neutrophil recruitment in murine HF. The chemotactic response of naive mouse neutrophils to plasma from naïve and HF mice (8 w post-MI) was measured by time-lapse confocal microscopy. ***A***, Representative tracks of DCFDA-loaded neutrophils (upper panels) and composite vector analysis showing rate and directedness of neutrophil migration (lower panels) towards plasma from naïve and HF mice, as well as in the absence of stimuli (no plasma control). Quantitation of distance traveled as line length (***B***), and rate of migration as line speed (***C***) for the three groups. Control n=106, naive n=198, HF n=217. ****p≤0.0001, ^##^p≤0.01, ^####^p≤0.0001 by two-tailed unpaired t-test. ***D***, Naïve mouse neutrophil transmigration in Transwell assays in response to control (no stimuli), G-CSF, CXCL1, CXCL5, mitochondrial formyl peptide, f-MNN, and control non-formylated peptide, MNN. No stimuli control n=9, G-CSF n=5, CXCL1 n=4, CXCL5 n=5, f-MNN n=5, MNN n=5. *p≤0.05, ****p≤0.0001, ^###^p≤0.001 by Kruskal-Wallis test with Dunn’s post-test (***B*** and ***C***) or one-way ANOVA with Dunnett’s post-test (***D***).

### Increased NETosis in murine HF

The biological effects of neutrophils depend in part on whether cellular cytotoxic functions are activated. NET production during neutrophil NETosis represents one important cytotoxic mechanism as the extracellular expulsion of nuclear DNA strands decorated with granular proteins and histones are strongly pro-inflammatory and tissue injurious.^16–18^ We assessed neutrophil NETosis and NET generation in both plasma and blood neutrophils isolated from mice with HF (8 w post-MI). Circulating plasma free NETs, indexed by either myeloperoxidase (MPO)-DNA or histone-DNA complexes (sandwich ELISA), were significantly increased in HF mice (*Figure 5A*). Next, MPO-DNA and histone-DNA complexes were measured by sandwich ELISA in neutrophils isolated from blood and maintained in culture for 4 h without stimulation. As illustrated in *Figure 5B*, both complexes were robustly increased in HF neutrophils as compared with naïve neutrophils. Histone citrullination, a consequence of peptidyl arginine deiminase-4 (PAD4) activation, is necessary for chromatin decondensation prior to the extracellular expulsion of NETotic nuclear DNA. Immunostaining of isolated neutrophils revealed colocalization of citrullinated histone H3 with total histones and MPO in expelled NETs by confocal microscopy, with significant increases of both total NETs and citrullinated NETs in HF neutrophils (*Figures 5C*). Taken together, these data establish augmented NETosis in ischemic cardiomyopathy and HF.

**Figure 5.**
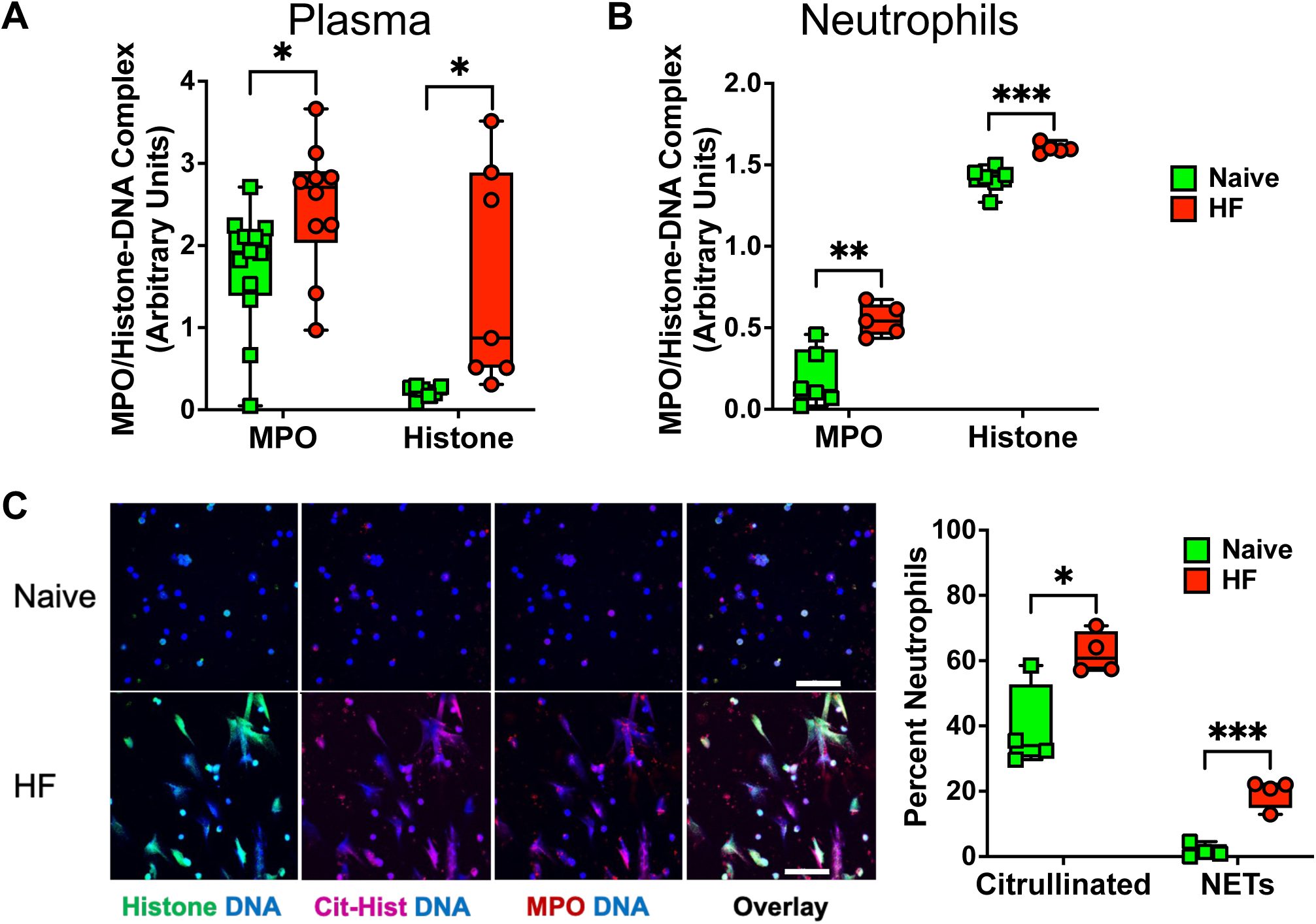
NETosis is increased in HF. Neutrophil NETosis was measured by myeloperoxidase (MPO)- and histone-DNA complex sandwich ELISA in plasma (***A***) and isolated, cultured peripheral blood neutrophils (***B***) from naïve and HF mice (8 w post-MI). Plasma MPO-DNA complex: naïve n=12, HF=10; Histone-DNA complex: naive n=7, HF n=7; Neutrophil MPO- and Histone-DNA complex: naïve n=6, HF n=5. ***C***, NET formation in HF was directly measured and quantitated in neutrophils isolated from naïve and HF mice by immunostaining of cultured cells for DNA (DAPI, blue), histone (green), citrullinated (Cit)-histone (magenta), and MPO (red). NETs were defined by co-localization of extracellular DNA with histone, cit-histone, and MPO by confocal microscopy. Representative immunofluorescence staining of naïve and HF neutrophils along with group data for percent of neutrophils staining for citrullinate histone alone and NETs are shown. Naive n=4, HF n=4; scale bar = 50 µm. *p≤0.05, **p≤0.01, ***p≤0.001 by two-tailed unpaired t-test.

### Neutrophil depletion abrogates progression of LV remodeling in HF

To determine the role of neutrophil expansion in ischemic cardiomyopathy, we performed 4 weeks of antibody-based neutrophil depletion initiated at two chronic HF timepoints – 4 w (intermediate stage) and 8 w (late stage) post-MI (*Figure 6A*). HF mice with comparable degrees of post-MI remodeling at these timepoints were randomly assigned to receive either anti-Ly6G neutralizing antibody (clone 1A8) or isotype IgG antibody (or vehicle) control, administered every 4 d for 4 more weeks. This dosing schedule was based on pilot studies demonstrating marked reductions in circulating neutrophils lasting at least 4 days with a single 1A8 dose (*<u>Figure S4A</u>*).

**Figure 6.**
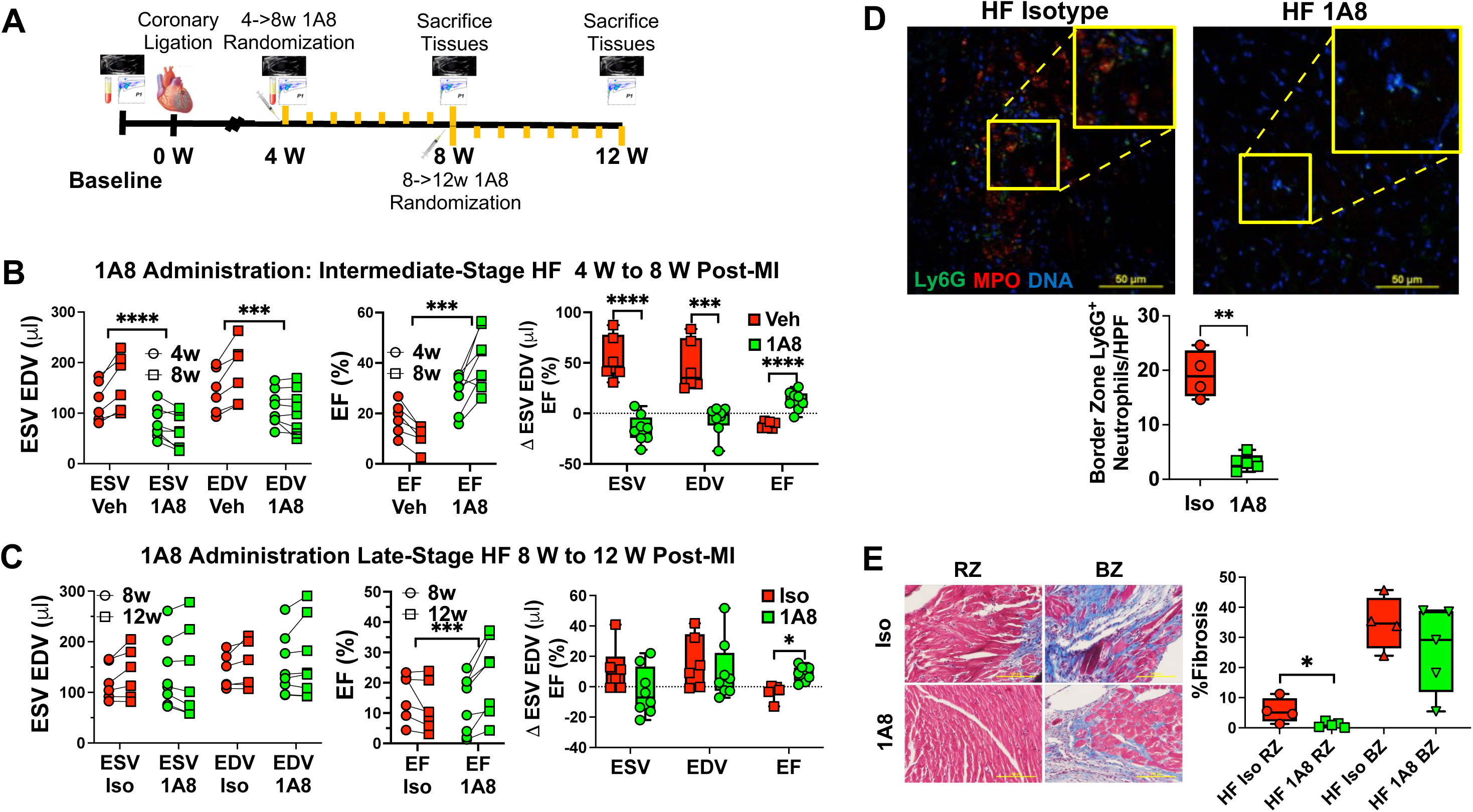
Neutrophil depletion alleviates LV dysfunction and remodeling in HF. *A*, Experimental schema of neutrophil depletion. Neutrophils were depleted in HF mice beginning at either 4 or 8 W post-MI with the anti-Ly6G neutralizing antibody, 1A8, or vehicle (Veh) or isotype (Iso) control (250 µg/injection/mouse i.p. q 4d). ***B*** and ***C***, Group data for absolute values and delta changes of LV end-systolic and end-diastolic volume (ESV and EDV) and ejection fraction (EF) by echocardiography, before and after antibody-based neutrophil depletion at an intermediate stage of HF (4 w to 8 w post-MI) (***B***) and late stage of HF (8 w to 12 w post-MI) (***C***). *p≤0.05, **p≤0.01, ***p≤0.001 relative to control by 2-Way ANOVA (ESV, EDV, and EF) or Mann-Whitney test (D ESV, EDV, EF); n number as indicated. ***D***, Immunofluorescent staining of isotype- and 1A8-treated late-stage HF hearts (border zone, 12 w post-MI) with Ly6G (green), MPO (red), and DNA (DAPI, blue). Iso n=4: 1A8 n=5. Scale bar 50 mm. **p≤0.01, two-tailed unpaired t-test.. ***E***, Representative Masson’s trichrome stains of isotype- and 1A8-treated late-stage HF hearts for evaluation of fibrosis in the remote zone (RZ) and border zone (BZ), together with fibrosis quantitation. HF isotype n=4; HF 1A8 n=5. Scale bar = 100 µm. *p≤0.05, two-tailed unpaired t-test.

HF mice at 4 w post-MI administered 1A8 for 4 weeks exhibited significant LVEF improvement, LVESV reduction, and LVEDV stabilization in contrast to vehicle-treated HF mice demonstrating the expected progressive worsening of these parameters (*Figure 6B*). Indeed, as compared to control HF mice, 1A8-treated HF mice exhibited significantly lower delta changes in ESV and EDV, and significantly higher delta EF over the 4-week treatment period. In the late-stage HF experiments, HF mice were randomized to 4-week treatment with either 1A8 or isotype IgG control beginning at 8 w post-MI. As illustrated in *Figure 6C and Figure S5*, while the 1A8-mediated effects on LV structure and function were less pronounced than with intermediate stage HF, neutrophil depletion even at this late stage resulted in less progression of LV remodeling, with smaller increases in chamber volume and significantly higher LVEF versus isotype-treated HF mice. Notably, 1A8 treatment profoundly decreased border zone cardiac neutrophil abundance in HF mice as assessed by immunostaining, establishing efficacy of cell depletion (*Figure 6D*). Along with lowering neutrophil number, 1A8 treatment markedly reduced cardiac interstitial fibrosis in HF mice, primarily in the remote zone, as compared to isotype IgG treatment (*Figure 6E*).

To complement the antibody-based neutrophil depletion, we performed analogous studies using genetic depletion of neutrophils in late-stage HF mice (8 w post-MI). For these we crossed Ly6G^Cre-tdTomato^ “Catchup” mice^21^ with ROSA26 Cre-inducible diphtheria toxin receptor (iDTR) mice that allowed neutrophil specific DTR expression (Ly6G-iDTR mice). Neutrophils were then depleted in Ly6G-iDTR HF mice via diphtheria toxin (DT)-induced apoptosis. Preliminary studies revealed a more modest ∼35% depletion of neutrophils after a single dose of 10 ng/g of DT given i.p., lasting approximately 5 days (*<u>Figure S4B</u>*). Hence, to ensure sufficient neutrophil depletion, DT was administered twice daily (ZT0 and ZT12) for 4 weeks (*<u>Figure S6A</u>*). As shown in *<u>Figure S6B</u>*, analogous to antibody-based depletion, genetic neutrophil depletion in Ly6G-iDTR late-stage HF mice resulted in attenuation of LV remodeling progression with stabilization of EDV and ESV, and better LV systolic function versus vehicle-treated Ly6G-iDTR HF mice. Collectively, these results establish an essential role for activated neutrophils in the progression of pathological LV remodeling in ischemic cardiomyopathy and HF.

### Increased stimuli for neutrophil trafficking in acute decompensated human HF

To explore clinical relevance, we next examined whether the milieu in human HF promotes neutrophil recruitment. Peripheral blood was collected from hospitalized patients with acute decompensated HF (ADHF) and ambulatory healthy controls. Select clinical characteristics of ADHF subjects are detailed in Table 2. The plasma was used to produce gradients for labeled naïve neutrophils, and chemotaxis quantified by time-lapse confocal imaging and vector analysis as described above. CXCL1, CXCL8, CCL3 chemokine gradients (baseline control without chemokines) were used in parallel as positive controls, given that these chemokines have been reported to be elevated in HF patients^36^ and also known to known to mediate neutrophil migration and recruitment. As illustrated in *Figure 7*, ADHF plasma provided a stronger chemotactic stimulus for naive neutrophils, as evidenced by significantly increased line length and line speed, and vector directedness as compared to normal control plasma. Moreover, the observed effect in ADHF was statistically comparable to the chemotaxis induced by supra-physiological levels of CXCL1, CXCL8, and CCL3. Hence, analogous to murine HF, the milieu in human HF is highly favorable for neutrophil trafficking.

**Figure 7.**
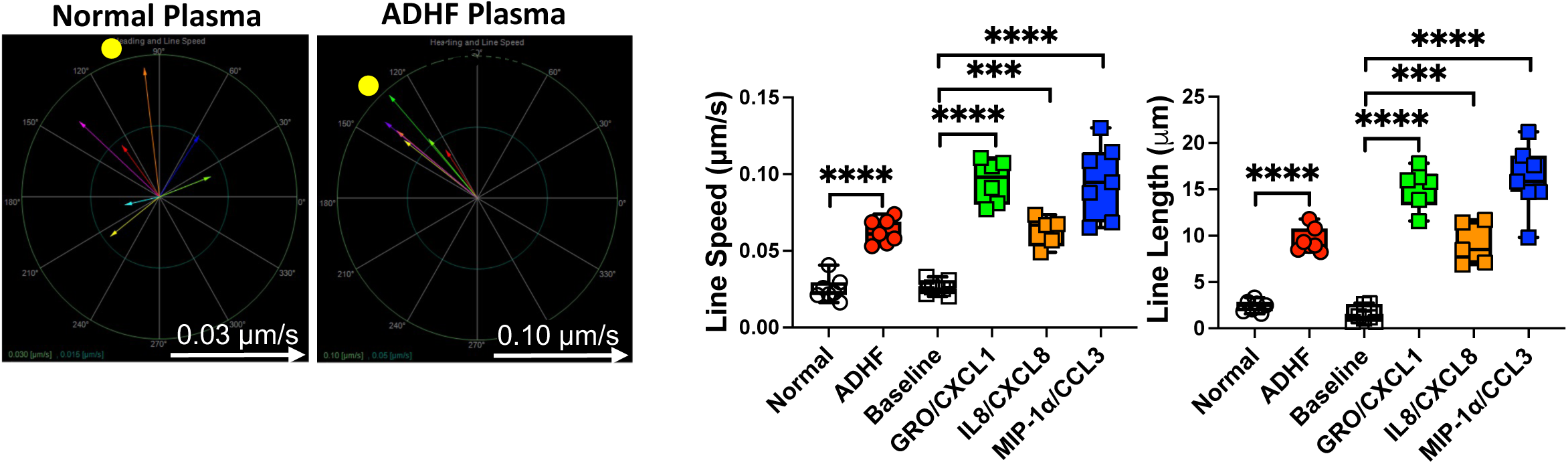
Human HF plasma promotes neutrophil chemotaxis. The chemotactic response of DCFDA-loaded naïve human neutrophils to plasma from normal and acute decompensated heart failure (ADHF) subjects was measured by time-lapse confocal microscopy. Representative composite vector analysis showing rate and directedness of neutrophil migration towards normal and ADHF plasma are shown, together with quantitation of rate of migration (line speed) and distance traveled (line length). Also shown for comparison are these parameters in response to gradients of single chemokines GRO/CXCL1, IL8/CXCL8 and MIP-1a/CCL3. Normal n=7, ADHF n=7, Baseline n=7, CXCL-1 n=6, CXCL-8 n=5, CCL-3 n=7. ***p≤0.001, ****p<0.0001 relative to control by one-way ANOVA with Dunnett’s post-test. Scale is 0.03 µm/s for control and 0.10 µm/s for ADHF.

**Table 2.** Select clinical characteristics of ADHF subjects.

| Parameter | ADHF subjects (n=7) |
| --- | --- |
| Age | 67 (53,82) |
| Female | 3 (43%) |
| Caucasian | 4 (57%) |
| LVEF | 27±5% |
| WBC (cells/μL) | 8,149±3,066 |

## DISCUSSION

We describe a heretofore unrecognized essential role for neutrophils in the progression of LV remodeling in *chronic* ischemic HF. There are several key findings. First, in chronic HF, neutrophils are expanded both locally in the failing myocardium (primarily in the border zone) and systemically in the blood, spleen and bone marrow together with increased BM granulopoiesis. Second, HF is characterized by heightened stimuli for neutrophil recruitment and trafficking with increased myocardial expression of CXCL1 and CXCL5 and increased neutrophil chemotactic factors in the circulation. Third, there is increased neutrophil NETosis in HF, evidenced by augmented neutrophil NET production and more circulating NETs. Fourth, neutrophil depletion with either antibody-based or genetic approaches abrogates the progression of LV remodeling at both intermediate and late stages of HF, with more pronounced effects in less advanced disease. Lastly, analogous to murine HF, the plasma milieu in human HF strongly promotes neutrophil trafficking. Taken together, these findings suggest that activated neutrophils and their associated cytotoxic products exert tissue-injurious effects in the failing heart that propagate adverse LV remodeling, fibrosis and dysfunction.

Following acute MI, neutrophils comprise the first wave of inflammatory cells that infiltrate the heart to clear dead cells and polarize macrophages toward a reparative phenotype, thereby ensuring fidelity of wound healing.^10, 11^ While their role in the initial phase of wound healing after MI is well established, whether neutrophil persistence and activation (analogous to macrophage expansion and recruitment^1, 4, 7, 37^) are important pathogenetic contributors to *chronic* LV remodeling and ischemic cardiomyopathy is poorly defined. Studies in human HF have demonstrated increased neutrophil lifespan that directly correlates with New York Heart Association (NYHA) class and LV systolic dysfunction.^38^ Plasma MPO and elastase, proteins secreted by activated neutrophils, are increased in patients with impaired left ventricular function and also correlate with the degree of dysfunction.^39^ Moreover, neutrophil counts predict the development of incident HF in subjects without cardiovascular disease,^40^ whereas the neutrophil to lymphocyte ratio predicts all-cause mortality in humans HF and in patients with ST elevation MI.^41, 42^ These studies suggest an important pathophysiological role for neutrophils in HF development and progression.

After MI and in HF, inflammatory hematopoiesis and myeloid cell production are increased in bone marrow and extramedullary sites such as the spleen due to a variety of factors, including increased sympathetic tone and reduced production of niche factors such as CXCL12.^37^ Consistent with this prior observation, in our study we demonstrate expansion of BM multipotent myeloid progenitors and GMPs in mice with HF, accompanied by persistently elevated neutrophil counts in the blood and the failing heart. While the expression of granulopoietic cytokines G-CSF and GM-CSF was not increased in the failing heart, there was higher expression of CXC chemokines CXCL1 and CXCL5, which together with augmented granulopoiesis and circulating neutrophils, establish conditions for increased transit and neutrophil recruitment in the myocardium.^33^ Notably, the plasma chemoattractant potential for neutrophils was also significantly increased in both murine and human HF, with a chemotactic driving force in human HF comparable to responses induced by classical neutrophil chemoattractants CXCL1, CXCL8, and CCL3. In murine HF, recruited neutrophils were most abundant in the MI border zone, potentially reflecting persistent preferential expression of neutrophil chemokines in residual viable areas most impacted by ischemic injury.^43^ Interestingly, plasma levels of CXC chemokines CXCL1, CXCL5, and CXCL8 are all increased in human HF, with CXCL8 levels correlating with NYHA class and LV dysfunction.^44^ Hence, combinatorial augmentation of neutrophil production, trafficking, and recruitment underly neutrophil expansion in the failing heart. This suggests the need for complementary therapeutic targeting of multiple convergent nodes to suppress neutrophil expansion in ischemic HF.

Persistent neutrophil tissue activity may lead to detrimental results in the heart via a variety of mechanisms. First, prior work has shown that direct adhesion of myeloid cells (*e.g.*, neutrophils and macrophages) to myocytes induces myocyte free radical generation and contractile dysfunction,^23, 45^ suggesting that direct cell to cell interaction of neutrophils and cardiomyocytes may contribute to contractile dysfunction in the failing heart. Indeed, the most pronounced effects of neutrophil depletion in our chronic ischemic HF model related to improvements in LVESV and LVEF, consistent with improved contractile function. Second, neutrophils and their secretory products serve to activate and recruit inflammatory monocytes.^17, 18^ Hence, neutrophil persistent in the failing heart may provide a favorable milieu for the observed expansion of pro-inflammatory infiltrating macrophages in the failing heart.^1, 4, 6, 7^ Third, neutrophils carry out toxic effector functions including the release of granule components and the formation and expulsion of NETs into the extracellular space during NETotic neutrophil cell death.^15, 16^ NETs are decorated with granular proteins (including neutrophil elastase and MPO), histones, mitochondrial formyl peptides and dsDNA which all can negatively impact cellular function. In our study we observed both increased circulating NETs and augmented NET release in circulating neutrophils in HF, suggesting that NETs may have a broad systemic impact in HF that could contribute to overall immune activation and myocardial remodeling. In line with this, studies of human HF have demonstrated increased plasma MPO that with NYHA functional class^46^ and parameters of adverse LV remodeling.^39^ Moreover, NETs, NET-specific markers (MPO/dsDNA), and NET-related tissue factor were increased in coronary plasma from the infarct-related artery and predictive of major adverse cardiac events (including cardiogenic shock and HF) in patients STEMI,^47^ consistent with a detrimental and cytotoxic impact on post-MI cardiac remodeling. Antibody-based and genetic deletion studies established an obligatory role for neutrophils in the development and progression of chronic ischemic HF. Both approaches attenuated the progression of LV remodeling, with greater therapeutic efficacy of neutrophil depletion when initiated at an earlier, intermediate stage (4 weeks post-MI) in heart failure. Interestingly, the most pronounced effect of neutrophil depletion on myocardial function was to stabilize (and not reverse) LV chamber size, but improve end-systolic volume and EF, suggesting a primary impact on contractile function during attenuation of LV remodeling progression as discussed above. These beneficial effects in chronic HF contrast with the detrimental effects of neutrophil depletion early after MI, which exacerbates post-MI remodeling related in part to loss of the beneficial effects of neutrophils on macrophage polarization in the context of wound healing.^10^ Hence, the biological effects of neutrophils in the post-MI heart is dependent upon disease stage, consistent with the idea of co-option of neutrophil behavior by tissues in accordance with their microenvironment and physiological demands.^48^ In this regard, non-ischemic mouse models of HF induced by pressure-overload^19^ and neurohormonal stress^20^ (angiotensin II [AngII] infusion) also exhibit cardiac neutrophil expansion early after induction of stress, but without persistence in the chronic phase as seen in ischemic cardiomyopathy. In these models, neutrophil depletion early in disease alleviated late cardiac remodeling. Interestingly, this beneficial effect was thought to be related to suppression of a KLF2-regulated NETosis pathway during AngII remodeling.^20^ Taken together, these observation underscore the need to carefully tailor therapeutic interventions targeting neutrophils to both the underlying etiology and disease stage of HF.

In summary, neutrophils are expanded locally and systemically in chronic ischemic HF, accompanied by heightened granulopoiesis, augmented stimuli for neutrophil trafficking and recruitment to the heart, and increased NETosis and NET formation. Targeted systemic neutrophil depletion at both intermediate and late stages of ischemic cardiomyopathy substantially reduces neutrophil abundance in the MI border zone and alleviates progression, but does not reverse, adverse LV remodeling while improving LV systolic function. Collectively, we demonstrate an essential role for neutrophil activation in the pathogenesis of ischemic cardiomyopathy and provide rationale for their therapeutic targeting in this disease.

