## Supplemental Data for "NEUTROPHILS ARE INDISPENSABLE FOR ADVERSE CARDIAC REMODELING IN HEART FAILURE"

[#]Equal author contribution

Correspondence to:
Sumanth D. Prabhu, MD
Professor and Chief
Division of Cardiology
Washington University School of Medicine
660 S. Euclid Ave, MSC 8086-43-13
St. Louis, MO 63110


or

Gregg Rokosh, PhD
Associate Professor
Division of Cardiology
Washington University School of Medicine
4565 McKinley Avenue, MSC-8086-12-780
St. Louis, MO 63110


#### SUPPLEMENTAL FIGURE LEGENDS

**Figure S1. Heart functional and morphometric indices in the chronic HF mouse model**. Functional, gravimetric, and phenotypic analysis confirmed HF 8 w after permanent coronary ligation. HF was verified in infarcted mice 8 w post-MI by demonstrated decreased EF (A), and LV end-systolic (ESV) and end-diastolic (EDV) dilation with representative parasternal long axis end-diastolic echocardiograms. (B) Sequelae to HF due to pulmonary congestion, systemic inflammation, and myocardial remodeling with heart, wet lung, and spleen weights, (Sham n=10, HF n=8) in sham and HF mice 8 weeks post-MI. (C) Fibrotic repair response to injury as degree of fibrosis in sham and remote and border zones of failing hearts assessed by Masson's trichrome staining with representative brightfield images. Scale bar = 100 μm. (H, I) Sham n=6, HF RZ n=6, HF BZ n=6. Significance denotes * p≤0.05, ** p≤0.01, *** p≤0.001, and **** p≤0.0001 relative to sham by two-tailed t-test (***A*** and ***B***) or one-way ANOVA with Dunnett's post-test (***C***).

**Figure S2. Representative neutrophil flow cytometry gating**. Neutrophils were defined as Singlet Cells CD45+CD11b+Ly6G+. Neutrophils were measured in (A) blood, (B) heart, and (C) spleen.

**Figure S3. Representative bone marrow flow cytometry hematopoietic progenitor and stem cell gating**. Bone marrow progenitors were defined as Lineage-Sca1-cKit+ (LKS-) and bone marrow stem cells were defined as Lineage-Sca1+cKit+ (LKS+). Progenitors were further broken down into granulocyte-monocyte progenitors defined as Lineage-Sca1-cKit+CD127-CD34+CD16/32+ (GMPs) and common myeloid progenitors defined as Lineage-Sca1-cKit+CD127-CD34+ CD16/32- (CMPs).

**Figure S4. Kinetics of antibody-based (1A8) and genetic diptheria toxin (DT)-based neutrophil ablation in mice.** Neutrophils were depleted in WT C57BL/6 mice with one dose of 250 µg 1A8 i.p. and in Ly6G-iDTR mice with one dose of 10 ng/g body weight DT. Neutrophils ($CD45^{+}CD11b^{+}Ly6G^{+}$ single cells) were quantitated daily by flow cytometry after 1A8 treatment (***A***) and DT treatment (***B***) to establish kinetics of depletion and repletion. 1A8 n=2, DT n=5.

**Figure S5.** Representative parasternal long axis echocardiograms of late-stage HF mice (8 w post-MI) receiving 4 weeks of treatment with either 1A8 or isotype IgG control. Echocardiography was performed on HF mice at baseline (***A***) prior to treatment (8 w post-MI) and after treatment (***B***) at 12 weeks post-MI. Representative images at end-systole are shown.

**Figure S6. Genetically targeted neutrophil depletion attenuates LV dysfunction in HF.** (***A***) Ly6G-iDTR HF mice (8 w post-MI) underwent neutrophil depletion with 10 ng/g DT or vehicle (PBS) administered at ZT0 and ZT12 for 4 weeks. ***B***, Group data for absolute values (Left and Middle) and delta changes (Right) for LV end-systolic and end-diastolic volume (ESV and EDV) and ejection fraction (EF) by echocardiography, before and after DT-based neutrophil depletion at late-stage HF, 8 w to 12 w post-MI. Vehicle n=5, DT n=12. Significance denotes * $p \leq 0.05$ by 2-Way ANOVA (ESV, EDV, and EF) and Mann-Whitney test (D ESV, EDV, EF).

### Figure S1

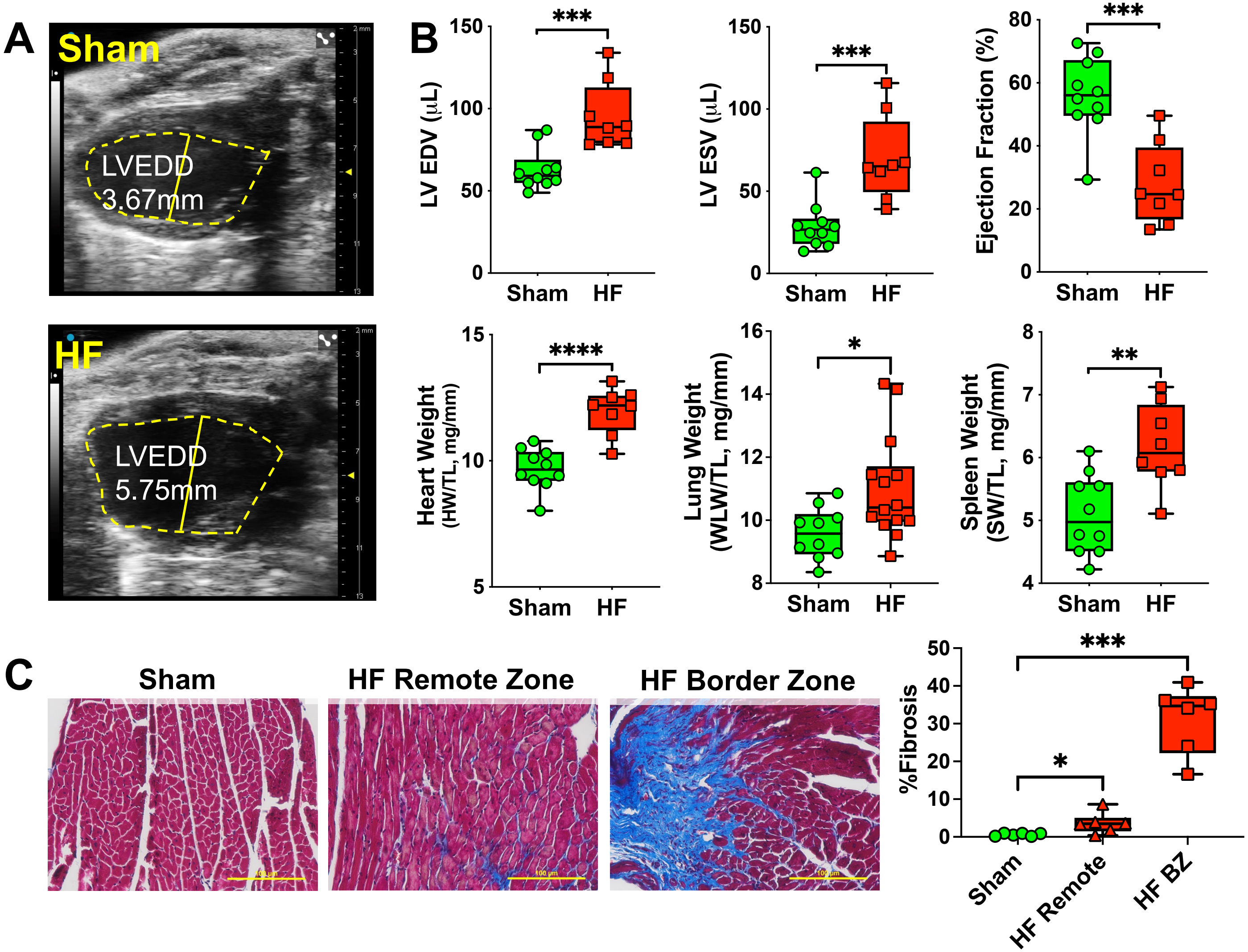


Figure S1. Heart functional and morphometric indices in the chronic HF mouse model. Functional, gravimetric, and phenotypic analysis confirmed HF 8 w after permanent coronary ligation. HF was verified in infarcted mice 8 w post-MI by demonstrated decreased EF (A), and LV end-systolic (ESV) and end-diastolic (EDV) dilation with representative parasternal long axis end-diastolic echocardiograms. (B) Sequelae to HF due to pulmonary congestion, systemic inflammation, and myocardial remodeling with heart, wet lung, and spleen weights, (Sham n=10, HF n=8) in sham and HF mice 8 weeks post-MI. (C) Fibrotic repair response to injury as degree of fibrosis in sham and remote and border zones of failing hearts assessed by Masson's trichrome staining with representative brightfield images. Scale bar = 100 μm. (H, I) Sham n=6, HF RZ n=6, HF BZ n=6. Significance denotes * p≤0.05, ** p≤0.01, *** p≤0.001, and **** p≤0.0001 relative to sham by two-tailed t-test (***A*** and ***B***) or one-way ANOVA with Dunnett's post-test (***C***).

Figure S2

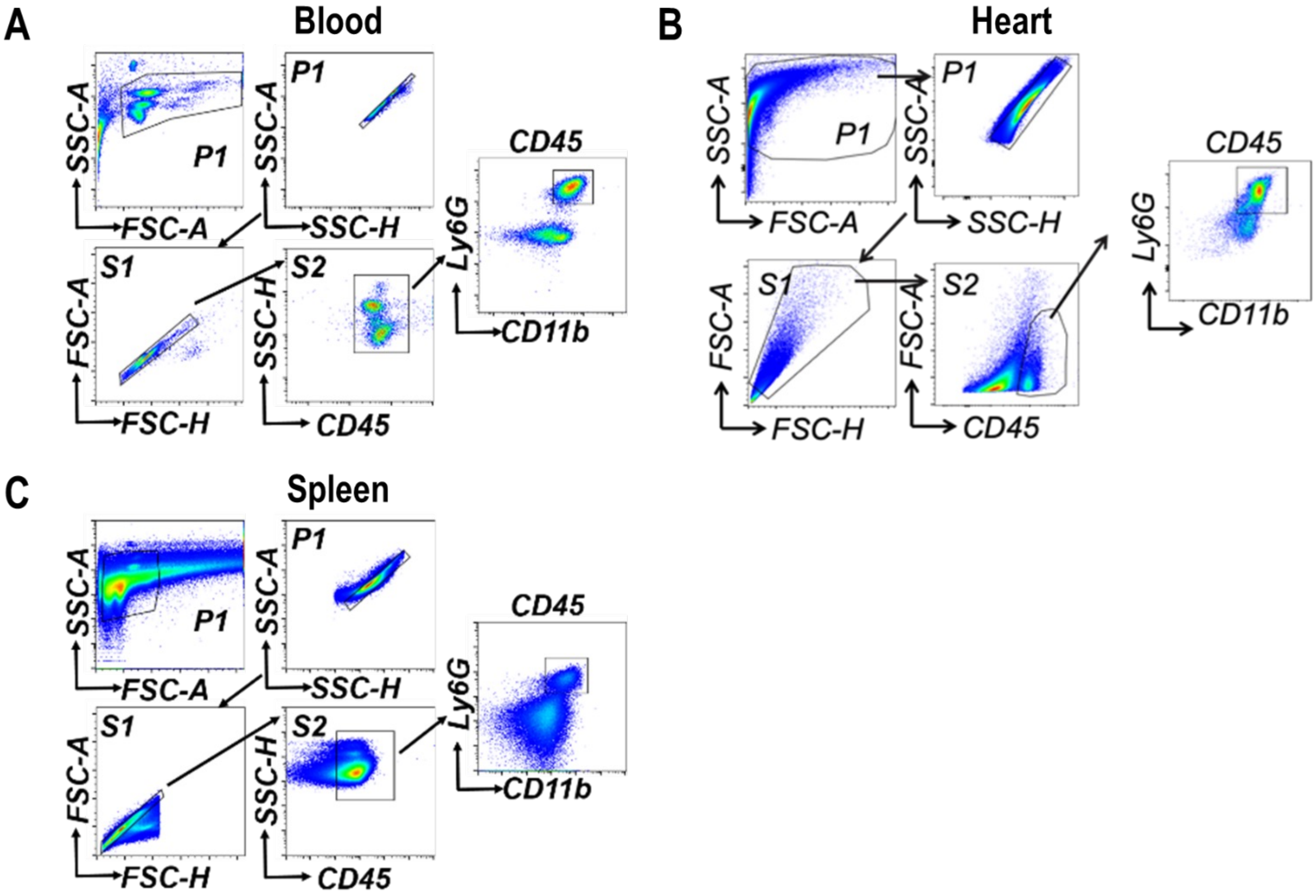


<u>Figure S2. Representative neutrophil flow cytometry gating.</u> Neutrophils were defined as Singlet Cells CD45+CD11b+Ly6G+. Neutrophils were measured in (A) blood, (B) heart, and (C) spleen.

### Figure S3

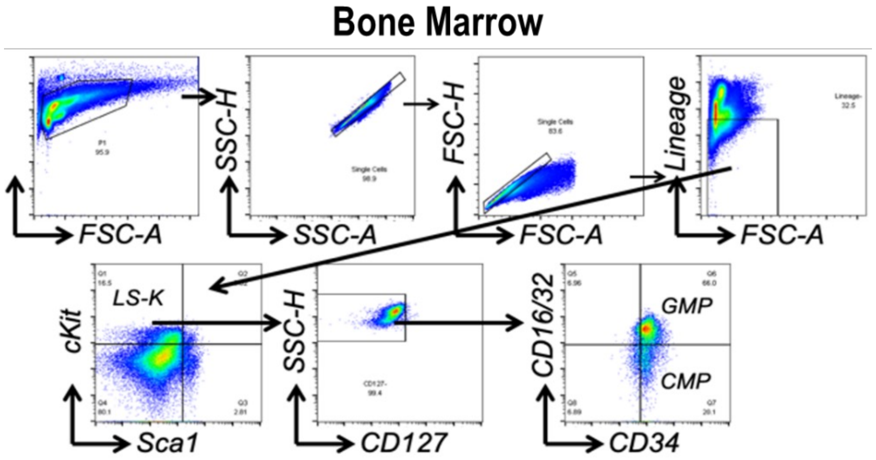


<u>Figure S3. Representative bone marrow flow cytometry hematopoietic progenitor and stem cell gating.</u> Bone marrow progenitors were defined as Lineage-Sca1-cKit+ (LKS-) and bone marrow stem cells were defined as Lineage-Sca1+cKit+ (LKS+). Progenitors were further broken down into granulocyte-monocyte progenitors defined as Lineage-Sca1-cKit+CD127-CD34+CD16/32+ (GMPs) and common myeloid progenitors defined as Lineage-Sca1-cKit+CD127-CD34+ CD16/32- (CMPs).

Figure S4

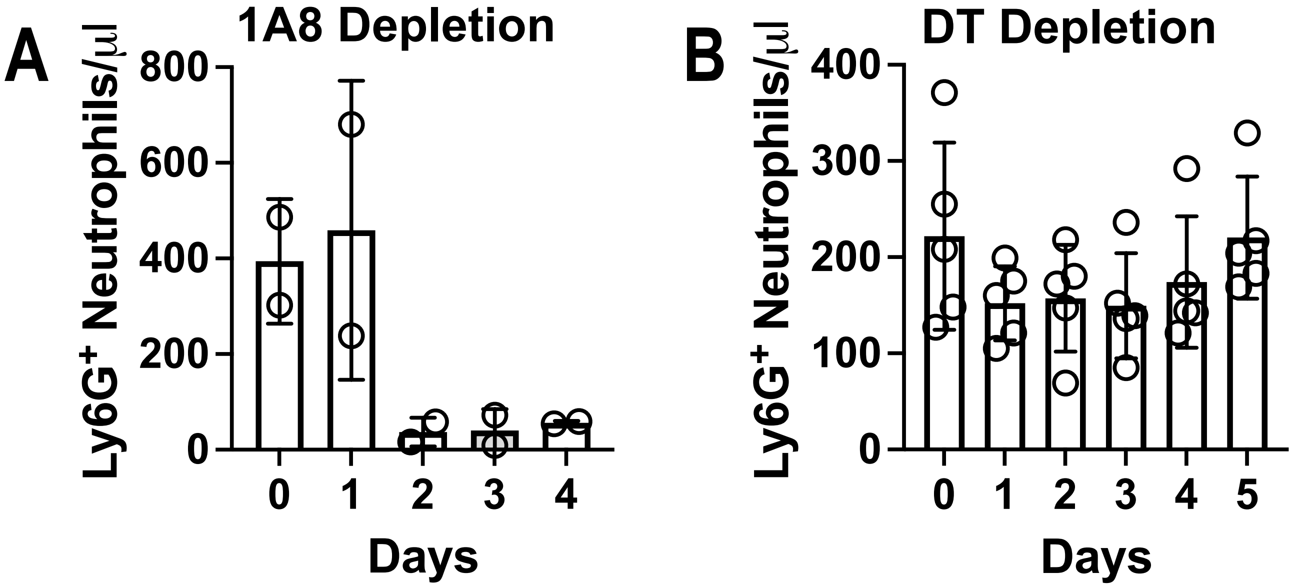


Figure S4. Kinetics of antibody-based (1A8) and genetic diptheria toxin (DT)-based neutrophil ablation in mice. Neutrophils were depleted in WT C57BL/6 mice with one dose of 250 μg 1A8 i.p. and in Ly6G-iDTR mice with one dose of 10 ng/g body weight DT. Neutrophils ($CD45^{+}CD11b^{+}Ly6G^{+}$ single cells ) were quantitated daily by flow cytometry after 1A8 treatment (***A***) and DT treatment (***B***) to establish kinetics of depletion and repletion. 1A8 n=2, DT n=5.

### Figure S5

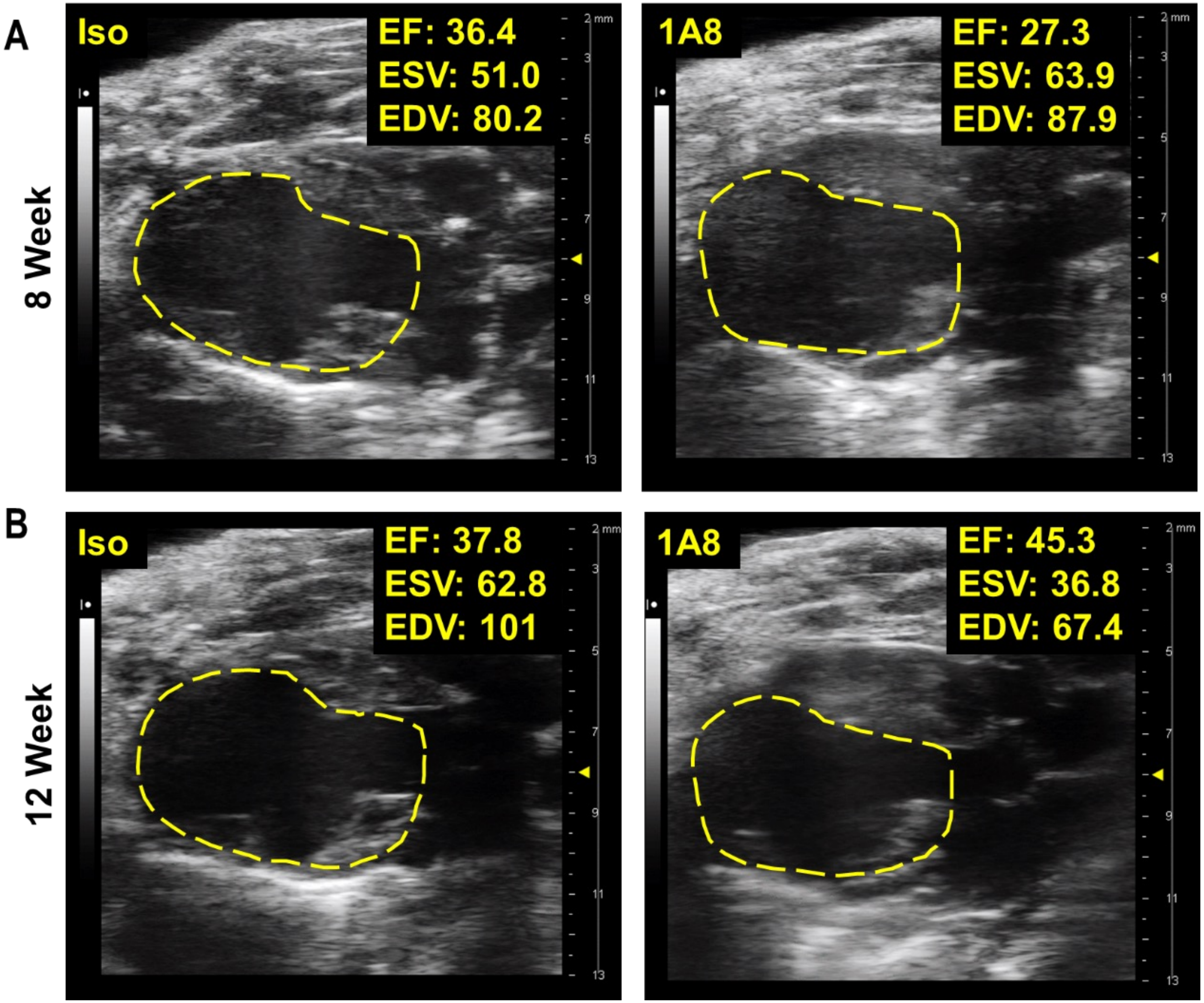


Figure S5. Representative parasternal long axis echocardiograms of late-stage HF mice (8 w post-MI) receiving 4 weeks of treatment with either 1A8 or isotype IgG control Echocardiography was performed on HF mice at baseline (***A***) prior to treatment (8 w post-MI) and after treatment (***B***) at 12 weeks post-MI. Representative images at end-systole are shown.

Figure S6

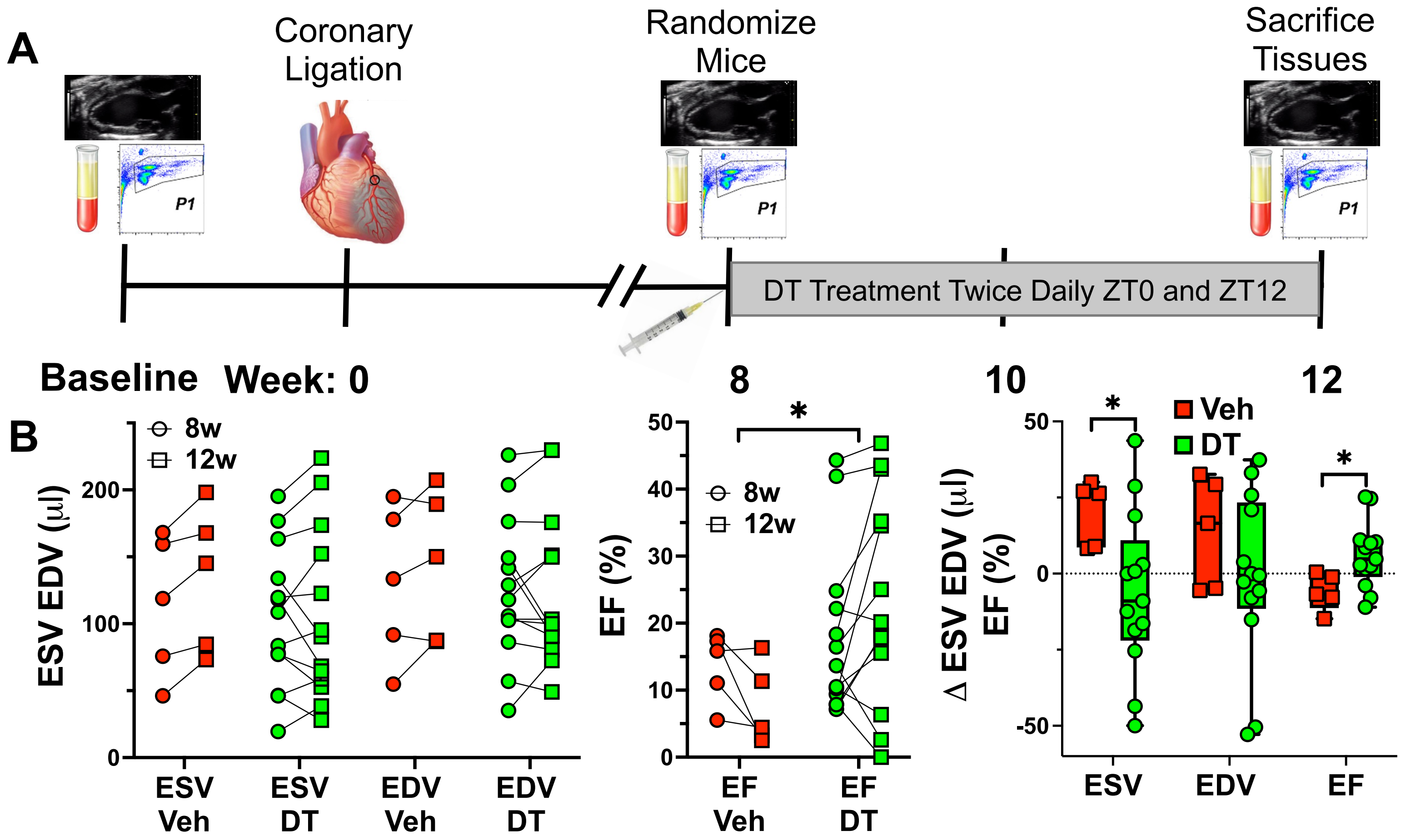


Figure S6. Genetically targeted neutrophil depletion attenuates LV dysfunction in HF. (***A***) Ly6G-iDTR HF mice (8 w post-MI) underwent neutrophil depletion with 10 ng/g DT or vehicle (PBS) administered at ZT0 and ZT12 for 4 weeks. ***B***, Group data for absolute values (Left and Middle) and delta changes (Right) for LV end-systolic and end-diastolic volume (ESV and EDV) and ejection fraction (EF) by echocardiography, before and after DT-based neutrophil depletion at late-stage HF, 8 w to 12 w post-MI. Vehicle n=5, DT n=12. Significance denotes * $p \leq 0.05$ by 2-Way ANOVA (ESV, EDV, and EF) and Mann-Whitney test (Δ ESV, EDV, EF).
